# Dynamical gene regulatory networks are tuned by transcriptional autoregulation with microRNA feedback

**DOI:** 10.1101/2020.02.03.932152

**Authors:** Thomas G Minchington, Sam Griffiths-Jones, Nancy Papalopulu

## Abstract

Concepts from dynamical systems theory, including multi-stability, oscillations, robustness and stochasticity, are increasingly implicated in the control of gene expression during cell fate decisions, inflammation and stem cell heterogeneity. However, the prevalence of the underlying structures within gene networks which drive these dynamical behaviours, such as direct autoregulation or feedback by microRNAs, is unknown.

We integrate transcription factor binding site (TFBS) and microRNA target data to generate a gene interaction network across 28 human tissues. This network was interrogated to identify network motifs capable of driving dynamical gene expression, in particular oscillations. Autoregulatory motifs were identified in 56% of transcription factors (TFs) investigated, 89% of which were also found in dual feedback motifs with a microRNA. Both the autoregulatory and dual feedback motifs were enriched in the network. TFs that autoregulate were found to be highly conserved between tissues. Dual feedback motifs with microRNAs were also conserved, but less so. Such dual feedback motifs were conserved between tissues, although TFs regulate different combinations of microRNAs in a tissue-dependent manner.

TFs which autoregulate are prevalent among human TFs and have more interactions with microRNAs than non-autoregulatory genes. The enrichment of such motifs within the human transcriptional network indicates that more genes may have interesting expression dynamics than previously thought. These data provide a resource for the identification of TFs which regulate the dynamical properties of human gene expression. These findings support the development of dynamical conceptual frameworks for the study of fundamental biological processes.

## Introduction

Cell fate changes are a key feature of development, regeneration and cancer, and are often thought of as a “landscape” that cells move through (Waddington, 1957; Huang *et al*., 2009). Cell fate changes are driven by changes in gene expression: turning genes on or off, or changing their levels above or below a threshold where a cell fate change occurs. “Omic” technologies have been successful in cataloguing changes in gene expression during cell fate transitions. Many computational tools have been developed for the ordering of gene expression changes in pseudotime, delineating cell fate bifurcation points and linking genes into networks (Aibar *et al*., 2017; Boukouvalas *et al*., 2018; Guo *et al*., 2019). However, while we have a good understanding of the fates/states that cells transition through and their order in time/space, the mechanisms that allow cells to move through the fate/state landscape are not well understood.

Gene regulatory networks are maps of interactions between different TFs, cofactors, and the genes or transcripts they target (Alon, 2007). Networks are commonly represented diagrammatically as graphs of the connecting components, such as TFs and their targets. Network motifs are small repeating patterns found within larger networks (Alon, 2007). Modelling of networks in this manner allows us to develop an understanding of how components interact and what behaviours they may generate (Alon, 2007; Neems and Kosak, 2010; Osella *et al*., 2011; Gennarino *et al*., 2012).

Although it is clear that gene interactions are dynamic, changing over time, current approaches in many biological studies focus on the qualitative analysis of genes or simple interactions between gene pairs: in short, how the perturbation of one gene affects the expression of another. However, gene expression is more nuanced than these traditional binary approaches can reveal. Biological networks are dynamical systems, that is systems that not only transform over time but resolve differentially depending on their parameter values, initial and boundary conditions, time delays, noise and the non-linearity of reactions. Autoregulation and cross-regulation of components are the heart of a dynamical network’s structure (Miller 2016). Dynamical systems are better suited than binary systems to explain biological phenomena because they can account for phenomena such as robustness and plasticity (DiFrisco and Jaeger, 2019).

In the conceptual framework of dynamical systems, oscillations can emerge as a ‘hallmark’ of a dynamical system that can transition into different attractor states. The fundamental properties that generate oscillations, such as the non-linearity of reactions, instability of components and time delays, are also the very properties that endow systems, including gene expression, with the ability to transition between different states. Oscillatory gene expression is now a well-recognised feature of several key transcription factors (TF)s and signalling molecules. For example, p53 is expressed in an oscillatory manner following DNA damage, leading to the arrest of the cell cycle and DNA-repair, although, sustained expression of p53 leads to cell senescence (reviewed in Hafner *et al*., 2019). p53 dynamics can also be altered in response to different stimuli (Batchelor *et al*., 2011). Oscillatory vs sustained expression of p53 may differentially regulate target genes, leading to various outcomes including cell cycle arrest, or growth and recovery (Purvis *et al*., 2012).

Another example is the oscillatory dynamics of the Hes genes, which play essential roles in many different developmental processes. Notably, oscillations in Hes7 govern the timing of the somite segmentation during embryogenesis (Bessho *et al*., 2003),, whereas oscillations of Hes1 have been shown to regulate the direction and timing of cell fate decisions in the developing neuronal system (Ohtsuka *et al*., 2001; Hatakeyama *et al*., 2004). Oscillations may be decoded by the phase, amplitude, or duration of the oscillatory phase (Batchelor *et al*., 2008; Levine *et al*., 2013; Sonnen *et al*., 2018).

While gene expression oscillations are important, they may be viewed as only one of the possible states that a gene network can assume. Dynamical systems can have non-intuitive behaviours, and it is necessary to employ mathematical modelling and quantitative approaches to understand their behaviour. Knowing which networks may show these properties is the first step. Coupling this approach to appropriate functional experimentation and mathematical modelling will allow a mechanistic understanding of cell fate/state transitions. Thus, we argue that discovering network motifs in biological networks that can drive a range of dynamics will facilitate a new conceptual and experimental approach to cell fate or cell state transitions, applicable to development, regeneration and disease, including cancer.

Results from synthetic biology show that the most straightforward motif that can produce oscillations is an autoregulatory negative feedback loop (Goodwin, 1965; Stricker *et al*., 2008). Autoregulation is a critical component of other oscillatory motifs, including the amplified feedback (Barkai and Leibler, 2000; Atkinson *et al*., 2003) and dual feedback loops (Stricker *et al*., 2008). A common element of the oscillatory motifs outlined here is negative feedback coupled with instability and delay. The repressilator, a synthetic gene interaction network of 3 repressors, forming a circuit of repression, utilised these principles to produce oscillatory expression of a fluorescent protein in Escherichia coli (Elowitz and Leibler, 2000).

In addition to the network structure, gene expression oscillations have other principles in common, such as time delays and instability of its components (mRNA and protein). Recent evidence suggests that mRNA stability is key to the generation of oscillatory gene expression (Moss Bendtsen *et al*., 2015), and this may be regulated by microRNAs. MicroRNAs are a class small non-coding RNAs around 22nts in length and are critical regulators of gene expression, modifying mRNA stability and translation (Kim, 2005). MicroRNAs are transcribed from either intergenic microRNA genes or intragenic loci producing primary transcripts containing hairpin loops. These primary transcripts are spliced into shorter pre-microRNA which are exported from the nucleus where they are subsequently processed into a double-stranded microRNA duplex (Ambros *et al*., 2003; Kim, 2005; Ha and Kim, 2014). One of these two transcripts, either the 5’ or 3’ arm, are selected to be the mature microRNA and enter into a repressor complex with Argonaute (AGO). MicroRNAs guide RISC (RNA-induced silencing complex) members to the 3’ UTRs of target messenger RNAs (mRNAs) (Kim, 2005; Ha and Kim, 2014). RISC protein AGO can then repress translation and increase the rate of mRNA deadenylation of target transcripts (Chen and Shyu, 2011). Increased deadenylation leads to decreased stability and therefore increased degradation of the target mRNA (Kim, 2005).

MicroRNAs are transcribed by Pol II (and in some cases Pol III) in the same way as mRNAs (Lee *et al*., 2004; Borchert *et al*., 2006; Monteys *et al*., 2010). As such, they are subject to the same regulatory input and are under the regulatory influence of transcriptional regulators such as TFs. MicroRNAs therefore, are often incorporated into biological gene regulation circuitry, such as in the case of the Hes1/miR-9 oscillator and p53 oscillatory modulation by miR-16 (Bonev *et al*., 2012; Issler and Mombach, 2017).

We have a good understanding of the structures that may allow dynamical behaviour in biological networks, such as bistability and oscillatory expression. However, the prevalence of such gene regulatory network structures is unknown. Are they common, or are they restricted to just a few cases of key TFs?

Here, to address this knowledge gap, we constructed a gene interaction network by integrating data from well characterised human TF and microRNA databases. We then interrogated it for the presence of network motifs that have the potential of dynamical gene expression, in particular oscillations. Specifically, we have used the previously identified network structures of known synthetic and natural periodically expressed genes to interrogate a set of human TFs for the presence of simple motifs that incorporate TF autoregulation and reciprocal interaction of microRNAs and TFs. We used well-annotated databases and drew on information for TF binding from ChIP-seq data incorporated from ReMap database (Chèneby *et al*., 2018), as well as microRNA target predictions from miRTarBase (Chou *et al*., 2018).

We report that motifs with the potential to generate oscillatory gene expression (hereafter termed “oscillatory motifs”) are widespread in TF networks. In particular, the autoregulatory motif, which is the “core” structure of all oscillatory motifs examined, is prevalent in our human TF dataset. We also demonstrate, through the use of tissue-specific sub-networks, that autoregulation with microRNA feedback is well conserved between tissues in terms of network topology. Surprisingly TFs were found to utilise different microRNAs for preserving the network structure in different cellular environments; possibly by utilising the expression of tissue-specific microRNAs.

## Results

### Most TFBS cluster in cis-regulatory modules

To understand the prevalence of network motifs within transcriptional and post-transcriptional networks, we collected datasets of experimentally-validated TF and microRNA interactions to construct a gene regulatory network. For the TFBS data, we took advantage of the ReMap project, which acquired human ChIP-seq experiments from GEO (Gene Expression Omnibus, Edgar *et al*. 2002). ReMap (v1.2) reprocessed the ChIP-seq data to identify 2,829 high-quality data sets (Griffon *et al*., 2015; Chèneby *et al*., 2018) and combined with ENCODE release V3 (Wang *et al*., 2013) TFBS data to annotate a total of 80,129,424 TFBS positions for 485 TFs.

The ReMap database defined a set of cis-regulatory modules, which are loci where more than one TF binds. The vast majority of binding sites fall within these defined cis-regulatory modules and have at least one overlapping peak from another TF (98.6%; Figure.1). We reasoned that TFBSs outside of these cis-regulatory modules were more likely to represent non-specific binding, and therefore excluded those sites, leaving 79,037,581 peaks, an average of 165,005 binding sites per TF.

**Figure 1:**
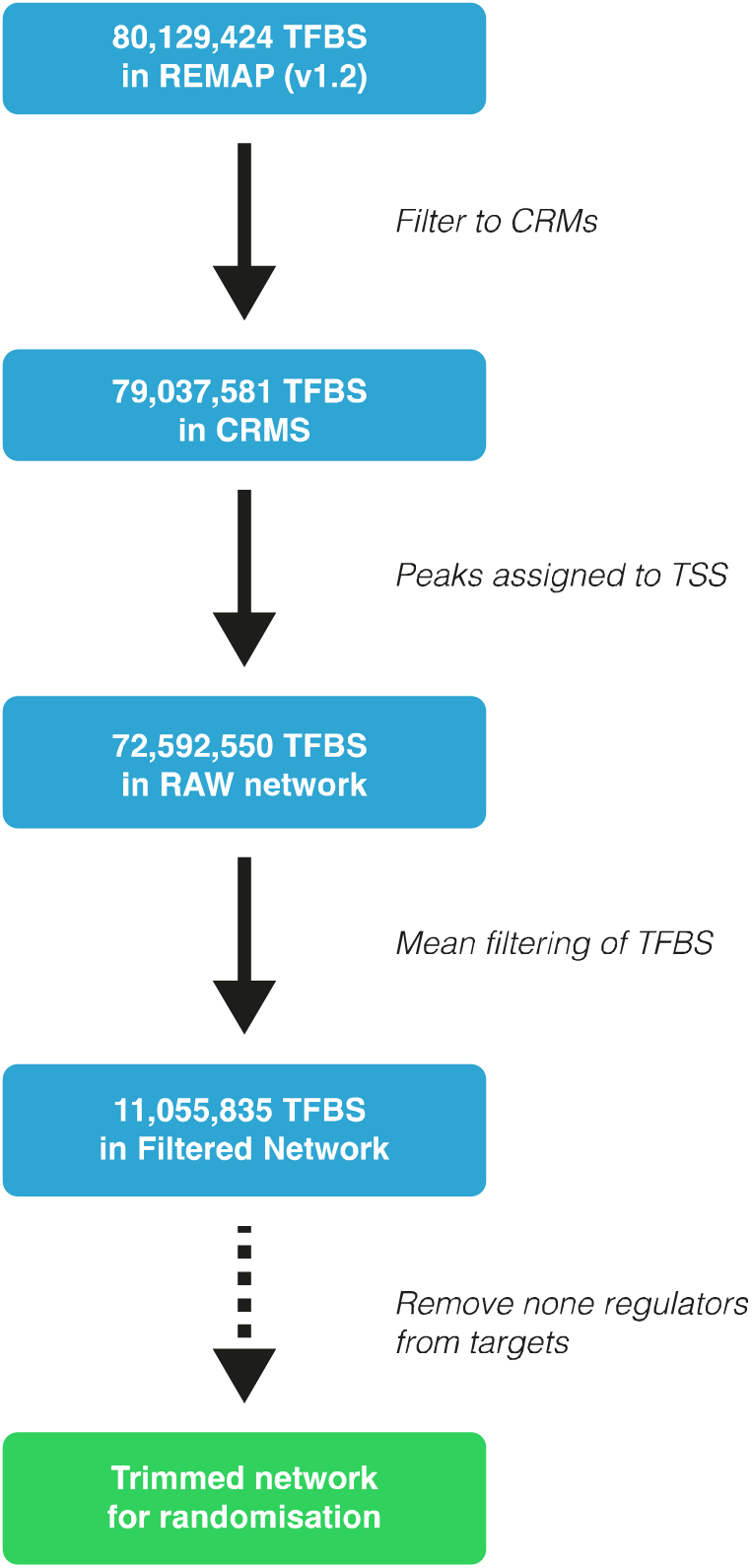
Refinement of TFBS data to final network. The original REMAP (v1.2) data contain 80,129,424 TFBS. These data were first filtered to include sites which were found in CRMs (regions where TFBS were found to overlap). These peaks were then assigned to TSS of all genes as annotated in Ensembl 89 in a window of 50kb upstream of the TSS and 10kb downstream of the TSS (*see methods*). To increase the confidence in the interactions between TFs and target genes, TSS-TF interaction were filtered based on the mean binding profiles of each TF (*see methods*). Interactions ≥ mean binding were maintained. This filtering produces the final network. For the randomisation experiments as all interactions in which we are interested contain feedback we produced a trimmed network which only contains genes which are both targets and regulators.

TFBSs were assigned to the closest annotated TSS in Ensembl within a range of 50kb upstream and 10kb downstream of the annotated TSS. This range is expected to contain the proximal promoter region of the gene, but may also include more proximal enhancers (Maston *et al*., 2006). We will therefore be discounting any transacting enhancers or silencers which fall outside of this range. Nonetheless, in the absence of chromatin accessibility information or DNA looping data, this relatively simple approach to the assignment of TFBS to genes should provide a good approximation of gene regulatory input. Of the 79,037,581 TFBS remaining after cis-regulatory module filtering, 72,592,550 TFBS (91.9%) were successfully assigned to TSSs of potential target genes (Figure. 1). The remaining 8% likely represent more distal enhancers and are not considered further here.

The set of annotated TSSs from Ensembl to which we have mapped our TFBS data include the 5’ ends of protein-coding transcripts, but also a number of classes of non-coding RNA, including lincRNAs and microRNAs. 68% of human microRNAs in our datasets overlap with longer transcript annotations; the majority of these microRNAs are found in introns of protein-coding genes. Intragenic microRNAs may be co-transcribed with their host genes or independently regulated (Baskerville and Bartel, 2005; Morlando *et al*., 2008). For example, Marsico *et al*., 2013 identified that around 60% of intragenic microRNAs share the transcriptional regulation of their host gene. To ensure that regulatory inputs for microRNAs are as complete as possible in our network, we combined the regulatory binding sites for the host gene with that of any TSS associated with the microRNA itself. On average across all intragenic microRNAs, 13% of the regulatory inputs in our network are associated with the microRNA alone, and 87% come from the host gene.

#### The raw network was completed by integrating microRNA target information

Next, we sought to integrate post-transcriptional regulation by microRNAs into our network. Several options are available for annotation of microRNA binding sites. For example, tools are available to predict microRNA target sites; most rely on seed matching (Bartel, 2009) that is, comparing the seed sequence of the microRNA with sequences within the 3’ UTRs of target genes. The signal-to-noise ratio for predictions using such short sequence matches is known to be relatively low, leading to high proportions of both false positives and false negatives (Oliveira *et al*., 2017). Instead, we collected microRNA target information from the miRTarBase (r7.0) database and integrated into the network. miRTarBase is an online database of validated microRNA targets, utilising experimental data from the literature, including luciferase reporter assays and CLIP-seq data (Chou *et al*., 2018).

### TFBS filtering selects for higher confidence interactions

Different TFs have substantially different numbers of targets in the raw network. For example, ZNF335 and MDM2 have one protein-coding target each while MYC and CTCF have over 18,500 targets, targeting over 93% of all protein-coding genes each. In the raw network, the median protein-coding target number for TFs as assigned is 10,651 (>53% of protein-coding genes).

More binding sites for a given TF at a given TSS has previously been shown to be a good indicator of regulation (Sikora-Wohlfeld *et al*., 2013). We therefore developed further filters to increase the certainty of the interactions, based on the distribution of number of binding sites for each TF across all TSSs. Essentially, we simply remove interactions between a TF and a TSS region where the number of binding sites for that TF is fewer than the mean number of binding sites per gene for that TF (*see methods*). In this way, we focus on TF/TSS region interactions where the number of TFBSs is above average. After this mean filtering is applied, the median number of protein-coding targets per TF is 4,303 (22% of protein-coding genes) a reduction of 60%. Some TFs are identified as targeting up to 9,369 protein-coding targets (47%, Figure.2A). Filtering TF-target interactions in this manner does not appear to bias or select for any particular target type, with the distribution of targets remaining proportional between different classes of gene targets, such as microRNA or proteincoding genes (Figure.2B).

**Figure 2:**
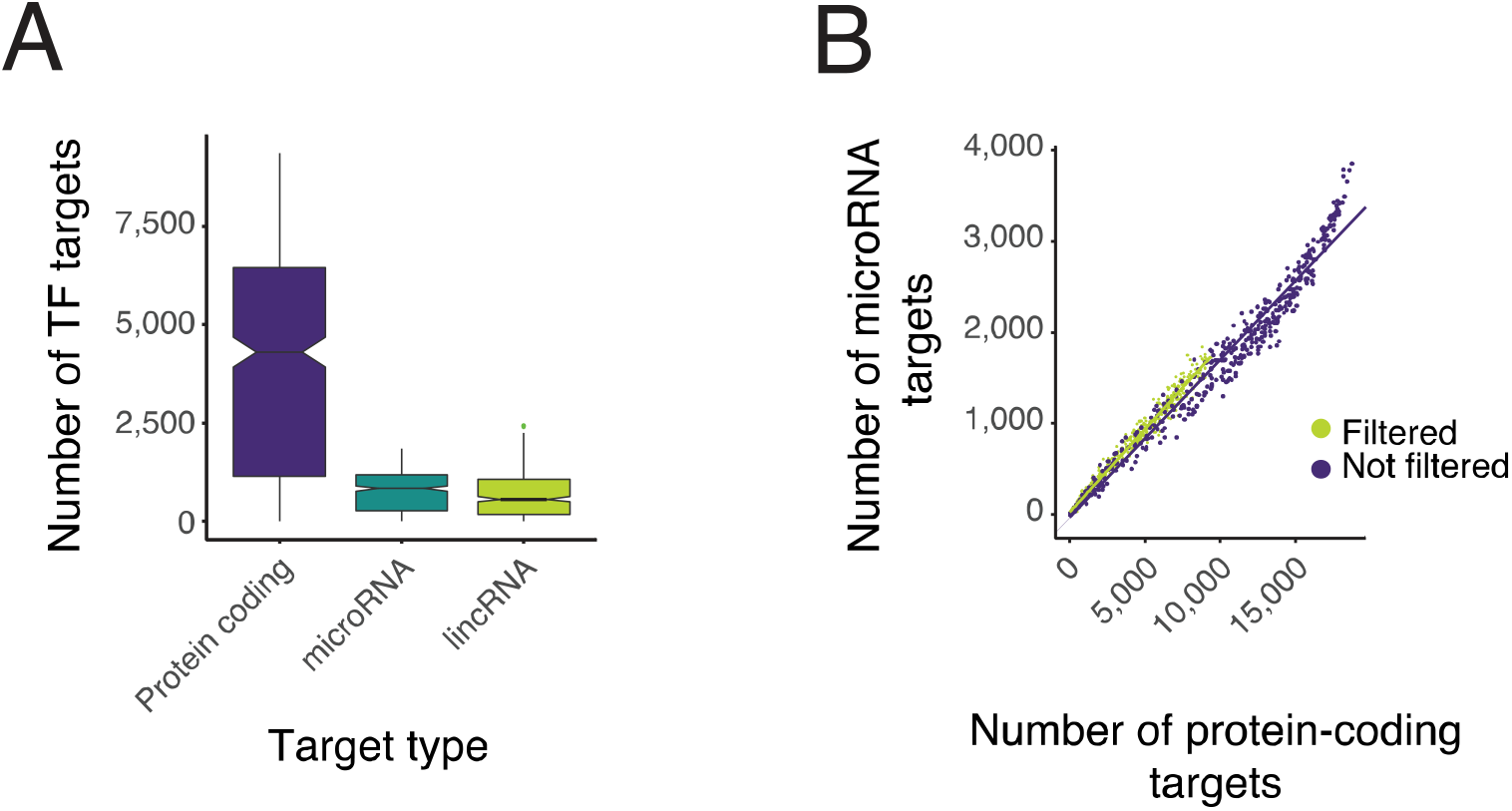
Network properties. (A) A boxplot showing the distribution of TF targets for protein coding, microRNA and lincRNA target types. (B) Shows the relationship between the number of protein-coding and microRNA targets per TF before and after filtering indicating that filtering of targets by our mean filtering method does not affect protein of microRNA targeting independently.

### The TF/microRNA regulatory network is highly connected

The complete network comprises 479 TFs (ReMap) and 2,599 microRNA (miRTarBase r7.0). The interactions in a network are described as edges that link one component (node) to another. These edges can be divided into two groups: inedges, which are receiving signals, such as a TF gene promoter receiving input from another TF; and out-edges, where a TF or microRNA regulates a promoter or mRNA. All components of a gene are linked together in our network. In the case of protein-coding genes, this means that the interactions at the TSS, mRNA and protein level all link to one component (node). The resulting network is highly connected: TFs on average have 120 in-edges and 170 out-edges, and microRNAs have 120 inedges and 145 out-edges. Since we are primarily interested in motifs where the targets of transcriptional and microRNA regulation are themselves regulators, we also produced a trimmed network using only edges that connect two regulators (TFs and microRNAs). In this trimmed network, the average TF protein has 200 in-edges and 830 out-edges, and microRNAs have an average of 120 in-edges and 7 out-edges. The difference here are due to the quantity of each type of network components, there are far fewer TFs in the network than microRNA.

### Transcriptional autoregulation with feedback by TFs and microRNAs is common

The network was interrogated to identify network motifs with structures that match those of existing oscillatory motifs. In all cases, the microRNA activity is assumed to be repressive, but the TF may be positively or negatively regulating. Nonetheless, we expect that a significant subset of the instances of the motifs identified will be connected in a way which could produce oscillations. The motifs selected were M1 (TF autoregulation), M2 (autoregulation with microRNA feedback), M3 (autoregulation with TF feedback), and M4 (two auto-regulators with co-regulation) (Figure.3A). TF autoregulation (M1) is a key feature of all our chosen network motifs, and was observed for 266 of 479 (56%) TFs in the network (Figure.3B). 237 of the autoregulating TFs (89%) are subject to microRNA feedback (M2). In total, the M2 motif was found 3,809 times within the network (Figure.3B), comprising combinations of 237 (49%) TFs and 1,254 (48%) microRNAs (Figure.3C). The M3 motif was observed 24,381 times and M4 19,622 times (Figure.3B). The TFs in M3 and M4 motifs have more targets than in the M2 motifs and are more highly interconnected (Figure.3D).

**Figure 3:**
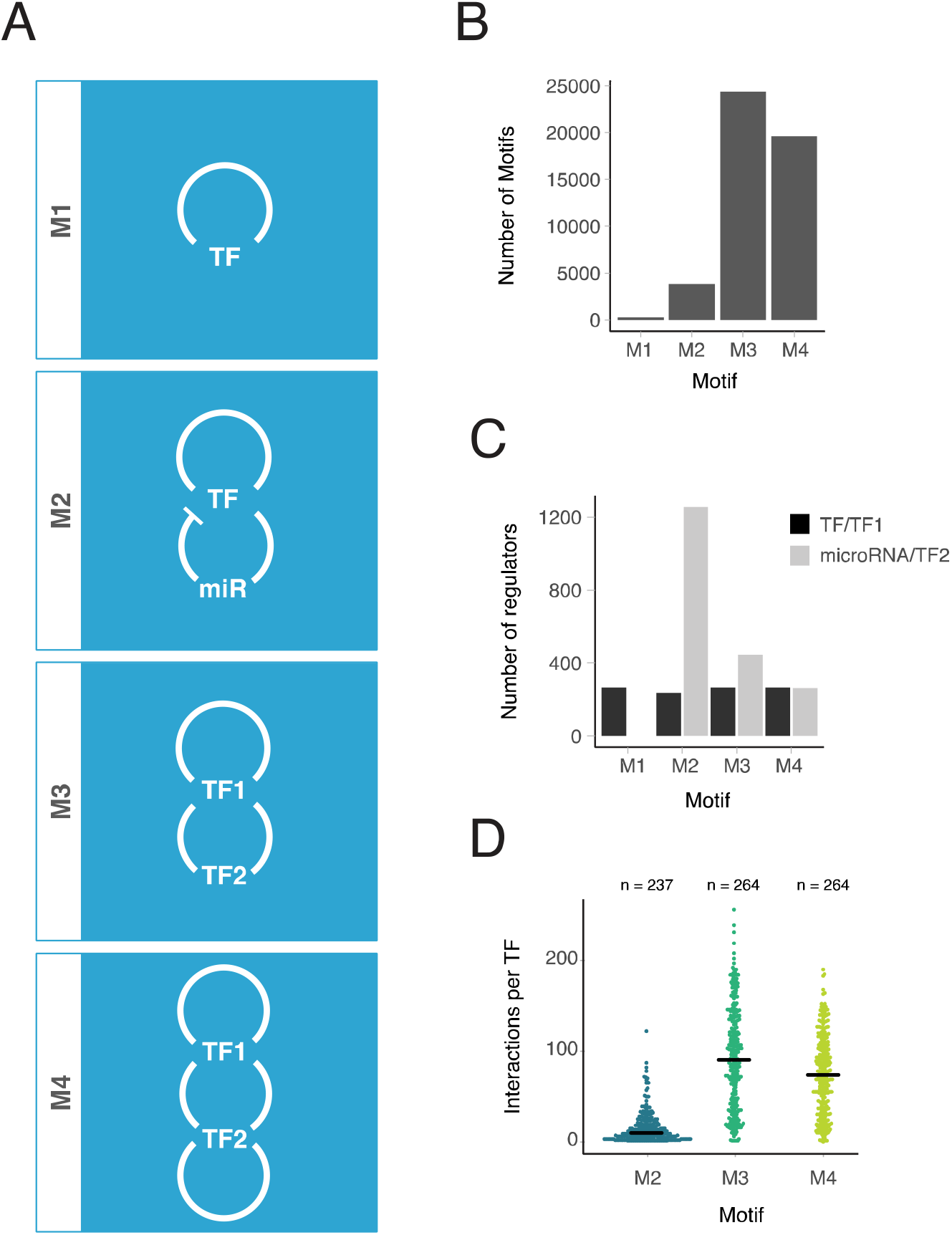
Network motifs are abundant in the gene interaction network. (A) A key to each motif shows each motif with the name below. (B) Bar graph showing the total number of motifs of each type within the network, M1 has the least at 266 though is a single component motif, M2 has 3,809 motifs, M3 is observed 24,381 times in the network and is the most abundant. M4 is observed 19,622 in the network. (C) Bar graph showing the number of TFs and microRNAs which contribute to each of the motifs. M2 motifs have the highest number of components due to the large number of. microRNAs in the network. (D) Beeswarm plot showing the number of other regulators each TF interacts within each motif. Black lines show medians. Reveals that while there are many more microRNAs in the network than TFs the TFs have more interconnections than the protein microRNA motifs.

A randomisation experiment was conducted to determine that the observed structure of the network is a product of underlying biology, not the inherent connectivity of the network. The trimmed network was randomised by rewiring while maintaining the node distributions of the network. All network motifs include an autoregulatory loop. If the autoregulatory loops (M1) are present more often than expected by chance, this could lead to all motifs appearing to be enriched. We therefore used two randomisation models: (1) “unlocked” randomisation, where all connections in the network are randomised. (2) “locked” randomisation, where autoregulatory edges were fixed. TF-target and microRNA-target interactions were randomised independently as they represent fundamentally different levels of control (transcriptional and post-transcriptional regulation, respectively). Separating the interaction types also acts to ensure the random networks retain structural comparability to the real network.

All motifs were found within the network more frequently than would be expected by chance. M1, M2 and M4 motifs were never observed in the 1000 random networks at numbers greater than or equal to their frequency within the real data (p < 0.001; M1 z=40.4; M2 z=6.10; M4 z=4.42). M3 network motifs were less significant than other motifs within the network, though they are still observed more frequently than would be expected by chance (p = 0.01; z=2.42). We therefore conclude that all four of the transcriptional autoregulation motifs, with and without feedback from TFs and microRNAs, are over-represented in the gene regulatory network. This overrepresentation is likely to reflect their importance in an underlying biological process.

### Autoregulatory genes are regulatory nodes in the network

It has been hypothesised that genes which autoregulate do so due to the need for more precise control over the expression of these genes (Crews and Pearson, 2009). Autoregulation, when negative, may also help buffer the response of the gene to multiple inputs to produce more robust expression behaviour. Connections into and out of autoregulatory genes were measured in comparison to non-autoregulatory genes to assess the connectivity of two groups in the network. Autoregulatory genes were found to have significantly more target genes than non-autoregulatory genes, an observation that holds across multiple target types (Figure.4A). The microRNAs within the network were also found to have significantly more autoregulatory targets than non-autoregulatory targets (Figure.4B). This observation did not appear to be specific to the group of microRNAs that show a preference for autoregulatory genes. Rather, most microRNAs (62%) regulate more autoregulatory genes than non-autoregulatory genes. Autoregulators on average have more inputs and outputs than non-auto-regulators independent of regulators or target type. These results taken together may indicate that auto-regulators form small local hubs in transcriptional and post-transcriptional networks. Tighter control of these genes, provided by autoregulation, may be necessary due to the high number of targets.

**Figure 4:**
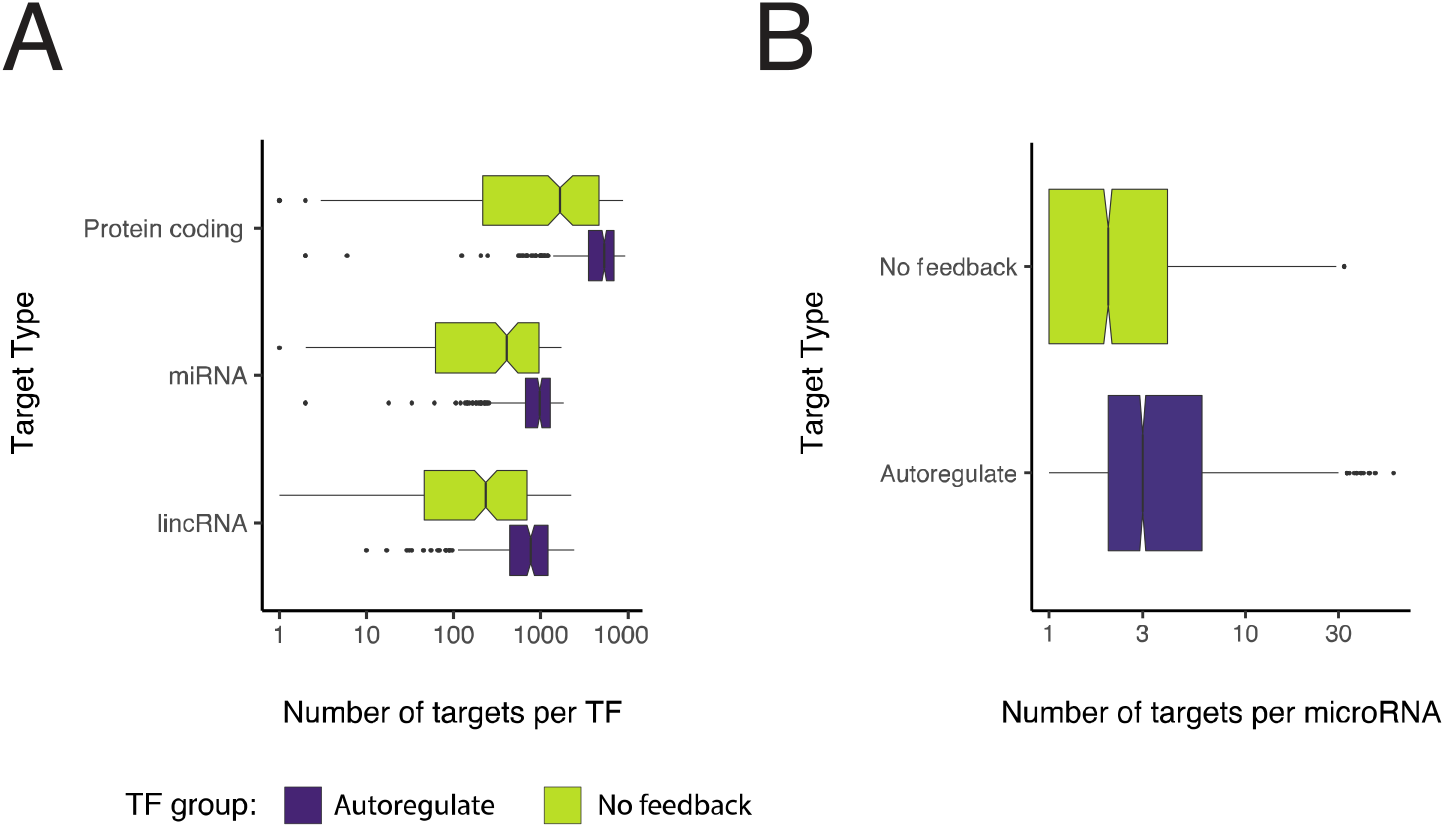
Autoregulatory genes are more connected than non-autoregulatory genes. (A) Boxplots showing the number, and gene type, of down-stream targets for both autoregulatory genes and non-autoregulatory genes reveal that autoregulatory genes have more targets than non-autoregulatory genes. (B) Shows the number of targets of microRNAs in the network which are autoregulatory or non-autoregulatory revealing that microRNA have more autoregulatory targets than non-autoregulatory targets.

### Tissue networks reveal conserved motifs between tissues

The ReMap data contains information on the cell types from which all TFBS data were obtained. We used this cell type information to group TFBS datasets into 28 tissue types. This information was exploited to identify patterns of interactions between TFs and microRNAs that were maintained between different tissue types. To this end, we built sets of tissue-specific networks involving TFs and microRNAs. Each tissue-specific network necessarily contains a subset of the nodes and edges present in the entire network. Of TFs which autoregulate, 43% had binding site data in only one tissue type, and were therefore not considered further here. 19% of autoregulating TFs were found in 2 tissues, 13% were observed in 3 tissues, and the remaining 24% were detected in four or more tissues (Figure.5A).

**Figure 5.**
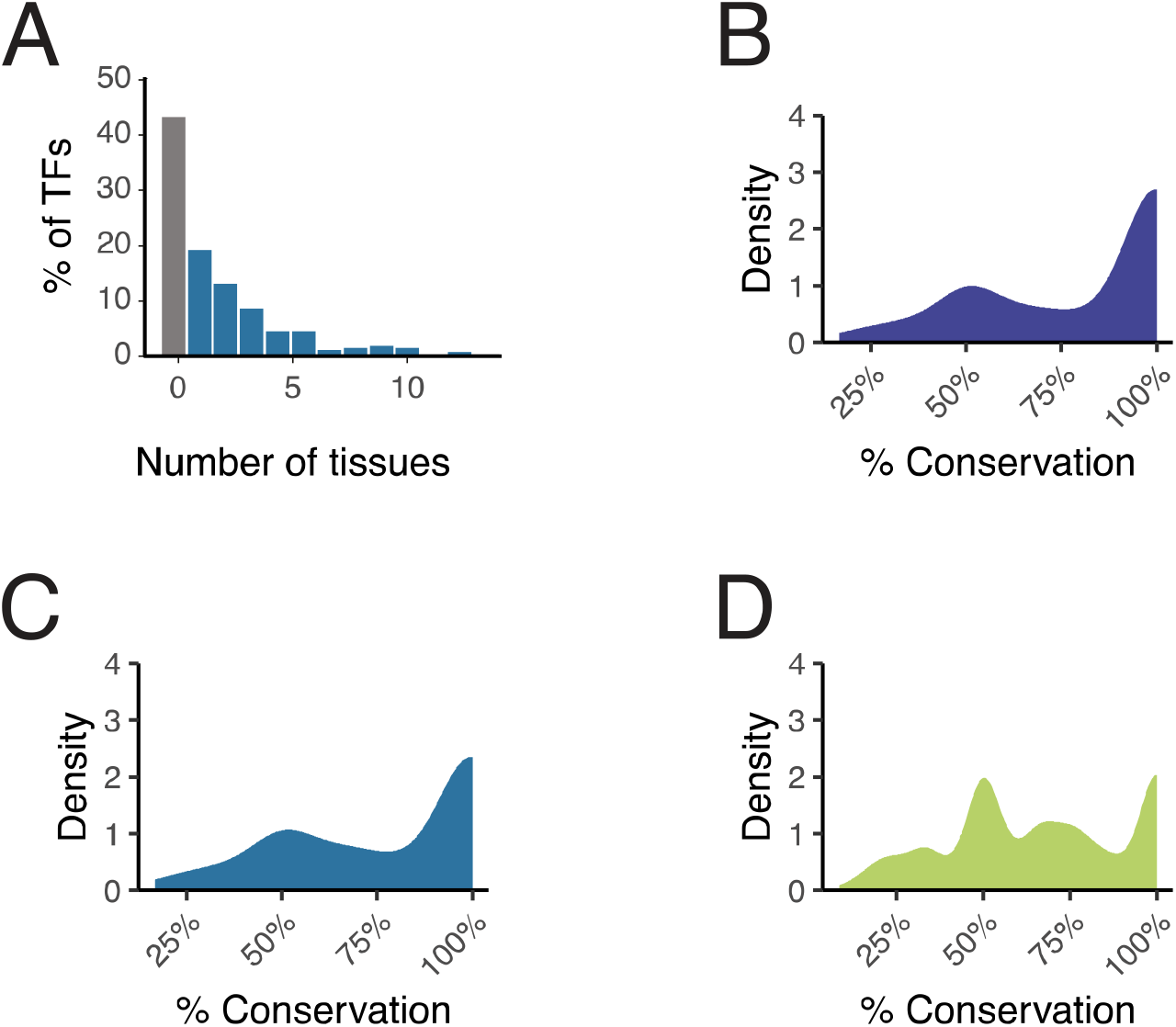
Conservation of TF in motifs between tissues. (A) Shows the percentage of TFs found in multiple tissues. 45% of TFs are limited to having data in only one tissue (grey bar) type with the majority having TFBS data in 2 or more tissues. (B) Distribution showing the conservation of M1 motifs across multiple tissues for TFs expressed in greater than one tissue. The majority of M1 motifs are highly conserved between tissues. (C) The conservation of M2 type motifs across multiple tissues for TFs expressed in greater than one tissue. (D) Shows the conservation of specific M2 type motifs with conserved microRNA-TF interactions, across multiple tissues for TFs expressed in greater than one tissue.

Where TFs have binding site data in more than one tissue, we asked whether their connections were conserved between those tissues. To this end, we investigated the conservation of M1 and M2 type motifs between tissues. M1 autoregulatory motifs are well conserved between different tissue types: the autoregulatory ability of over half of all TFs was conserved in 100% of tissues where data exist (Figure.5B). This suggests that their biological function may be linked to their ability to autoregulate. The presence of M2 motifs was only slightly less well conserved between tissues (Figure.5C). When M2 motifs are not conserved between tissues, the most significant factor is loss of autoregulation – 19% of all M2 motifs lose their autoregulatory signal between tissues, whereas only 1.3% of motifs are not conserved because they are missing TF regulation of the microRNA. However, even when the presence of the M2 motif is conserved, the identity of the components of the motifs may change. 18% of autoregulating TFs in M2 motifs, while retaining a core set of microRNAs, target different miRNAs in different tissues. The data suggest that the specific combinations of microRNAs and TFs vary between tissues, but the M2 motif structures themselves are maintained, sometimes by co-option of different microRNAs into those regulatory processes (Figure.5D).

Overall, our data show that autoregulation of TFs is prevalent and that microRNAs engage more frequently with autoregulated TFs. Indirect feedback, both with microRNAs and between TFs is also widespread within in the network. The variety and quantity of feedback observed here, coupled with the conservation for these network structures between tissues, may indicate an important role for these motifs in conferring dynamic behaviour to the TFs involved.

## Discussion

We generated a gene interaction network containing both transcriptional and posttranscriptional regulation. Previous work has indicated the high prevalence of autoregulatory loops in the human gene regulation network (Kiełbasa and Vingron, 2008). Kiełbasa and Vingron (2008) used computational TF target predictions based on binding DNA motifs to look for autoregulatory loops. Here we build upon this work by utilising experimentally predicted gene targets, rather than relying on DNA binding motifs, while also increasing the number of genes investigated from 292 TFs to 479. Our data also include the addition of post transcriptional feedback with the inclusion of microRNA target data. The motifs investigated in this study are capable of producing a range of different dynamic behaviours in their component genes, including noise modulation, bistability and oscillatory expression. Analysing the prevalence and behaviour of these motifs is therefore essential to our understanding of fundamental gene regulatory processes and their resulting molecular and cellular phenotypes.

Autoregulating TFs are highly enriched within our network, consistent with previous observations (Thieffry *et al*., 1998; Kiełbasa and Vingron, 2008). These autoregulatory genes are also highly connected within the network, possessing more inputs and outputs to protein and microRNA genes than non-autoregulatory genes. The high number of connections of autoregulatory genes indicates that they play central roles in processing both inputs and outputs in the network. The wide range of gene expression dynamics which can arise from autoregulation may explain the prevalence of the motif in biological networks.

Autoregulation (M1, Figure.3A) can produce many different behaviours depending on the properties of the components involved, and the type of inputs into the system. Negative autoregulation has previously been shown to decrease variability in the expression of a gene, buffering the noise of transcription and translation (Becskel and Serrano, 2000). Negative autoregulation can also decrease the variability in the levels of a gene in a population of cells. Conversely, positive autoregulation can increase variability of protein levels between cells in a population leading to bistability (Becskei *et al*., 2001). Negative autoregulation may also decrease the response time of a gene when compared with simple regulation of one gene targeting another (Rosenfeld *et al*., 2002). The response time of a gene is the time required to achieve 50% the concentration of steady-state (Alon, 2007). Steady-state is achieved when the degradation and production rates of the gene products are balanced. Negative autoregulation adjusts the balance between degradation and production by decreasing the transcription rate as the level of protein increases. Therefore negative autoregulation leads to faster equilibrium, as the degradation rate matches the production rate earlier. (Rosenfeld *et al*., 2002). Positive autoregulation has the opposite effect: the rate of transcription increases as the levels of protein increase, and the system therefore takes longer to reach an equilibrium state (Maeda and Sano, 2006; Mitrophanov *et al*., 2010).

MicroRNA feedback has also been previously predicted to be an enriched motif in human and mouse gene networks (Tsang *et al*., 2007). While microRNAs themselves are often suggested to act as buffers of gene expression, the interaction between a repressing TF and a microRNA in an M2 motif under certain conditions may act as a bistable switch (Goodfellow *et al*., 2014). For example, ZEB and miR-200 form a negative feedback loop where miR-200 inhibits mZEB and ZEB inhibits the transcription of miR-200. This loop is thought to be involved in mesenchymal to epithelial-mesenchymal transition (Lamouille *et al*., 2013). Modelling of miR-200 and ZEB has shown the system to switch states depending on input: high levels of miR-200 and low ZEB levels favour the epithelial phenotype, while low miR-200 low and high ZEB produce a mesenchymal phenotype (Tian *et al*., 2013; Lai *et al*., 2016). The M2 motif involving ZEB1 and miR-200b-3p is present in our network analysis (S2).

One of the goals of this work was to identify motifs with similar structures to existing oscillators, and thereby predict new oscillators. Every motif investigated in this study is capable of producing oscillatory patterns of gene expression, given the correct properties of the component genes and gene products. Oscillatory gene expression is an emerging area of research across many fields of biology including inflammation, stem cell heterogeneity and neurogenesis (Graf and Stadtfeld, 2008; Kobayashi and Kageyama, 2010; Nguyen *et al*., 2013; Shimojo and Kageyama, 2016). The core conditions required to produce oscillations are negative feedback, instability and delay (Monk, 2003; Bratsun *et al*., 2005; Novák and Tyson, 2008). Negative autoregulation is the most straightforward oscillatory motif and was the first synthetic oscillator to be described (Goodwin, 1965). Oscillations can occur in just this simple system if a gene’s mRNA and protein half-lives are shorter than the delay between gene activation and autorepression (Novák and Tyson, 2008). Negative autoregulation was shown to produce noisy oscillations in vivo when implemented through a synthetic autoregulatory module (Stricker *et al*., 2008).

The addition of negative feedback with a microRNA modifies the M1 motif to the M2 motif. This motif matches the network structure of the known ultradian oscillator HES1, a highly conserved oscillatory gene which drives cell fate decisions during neuron genesis (Ohtsuka *et al*., 2001; Hirata *et al*., 2002). HES1 partners miR-9 in an M2 motif, and the levels of miR-9 are able to drive neuronal differentiation through the modulation of HES1 dynamics (Bonev *et al*., 2012; Tan *et al*., 2012). It has also been hypothesised that the microRNA may also act as a self-limiting timer accumulating gradually through rounds of oscillations to a level where it destabilises the oscillations (Bonev *et al*., 2012).

The M3 motif (Figure.3A) is similar in structure to the M2 motif, with the microRNA substituted for a TF. In cases where the autoregulator in the M3 motif is a positive regulator of its own transcription, and the second TF negatively regulates the first, the M3 motif has been shown in synthetic biology to generate oscillations in vivo (Atkinson *et al*., 2003). The M4 motif is structurally similar to the M3 motif; however, both TFs are autoregulatory. Work by Stricker *et al*., 2008 has shown that the M4 motif can produce robust oscillations in situations where one TF is a positive regulator of transcription, and the other is negative.

Instances of autoregulation (M1) and autoregulation with microRNA feedback (M2) are highly conserved between tissues. The level of conservation indicates that these network motifs are necessary for core functional roles in different tissues. The component TFs were found to be conserved in M2 motifs in multiple tissues, but in many cases regulating different microRNAs between different tissues. Previous studies have shown that microRNAs have highly specific patterns of tissue expression (Ludwig *et al*., 2016). The switching of microRNAs in M2 motifs between tissues may therefore be due to differential incorporation of microRNAs into the network based on their availability in those tissues.

Our data show that autoregulation is prevalent amongst TFs and that genes which autoregulate are highly integrated into transcriptional networks. Further, autoregulatory genes preferentially engage with microRNAs, targeting and targeted by more microRNA than non-autoregulatory genes. All of the oscillatory type motifs were identified more than would be expected by chance within our data. The wide range of behaviours which TFs can exhibit when contained within these motifs suggest that these network structures of TFs, microRNAs and their target genes allow dynamic expression to control core cellular processes. The type of short time scale unsynchronised dynamics that these motifs predict is difficult to detect experimentally without real time single cell analysis. Our data therefore provides an invaluable resource in the search for dynamically expressed genes.

## Methods

All scripts can be found at https://github.com/TMinchington/network_codes. All graphs were generated using R (R Core Team, 2019) and the ggplot2 library (Wickham, 2016).

### ReMap data filtering

TFBS data were obtained from the ReMap2018 (v1.2) database (hg38, All peaks). The cis-regulatory module data were also downloaded from ReMap2018 for the matching version. ReMap2018 (v1.2) contains TFBS data for 485 TFs. TF datasets generated using non-specific antibodies were removed from the data. For example, the binding data for RUNX contains bindings which are non-specific and may relate to any member of the RUNX family. We removed tracks were where non-specificity was evident from the ReMap annotation. After this filtering was applied, 479 TFs remained. Transcriptional regulator binding sites were filtered to remove sites which were not found within cis-regulatory modules as described in the ReMap paper (Chèneby *et al*., 2018). The filtering was performed by running script chip_in_cmfs.py (arguments: CRM_file, chip_file). This custom python script uses the coordinates of the cis-regulatory modules from the ReMap database to cycle through the ReMap TFBS file and outputs TFBS which have loci contained within the cis-regulatory modules based on the genome coordinates.

### Assigning TFBSs to target genes

Human gene annotation data were downloaded from Ensembl 89 BioMart (human genes; GRCh38.p10). TFBS binding site coordinates for 485 TFs were utilised from the ReMap database (V1.2) (Chèneby et al. 2018). TFBS (ReMap (v1.2), Chèneby et al. (2018)) were assigned to the TSS (Ensembl 89) which was closest within a 60kb region (10kb downstream and 50kb upstream of TSSs) using the custom python scripts peak_MULTI.py and peakME_functions.py.

### Assigning microRNAs to host transcripts

The coordinates of human genes and exons were downloaded from Ensembl (89, GRCh38.p10, Yates *et al*., 2016). The genomic positions of microRNAs were downloaded from miRBase FTP server (release 22.0, Kozomara and Griffiths-Jones, 2011). MicroRNAs were then assigned to all transcripts with which the annotations overlap. Overlap with transcripts and positions within the transcripts were performed using microME.py. MicroRNAs within the peak_MULTI.py output files then inherited a copy of the TSS regulation of the genes in which they were contained while also maintaining the TSS as listed in the Ensembl 89 data.

### Generating the transcriptional and translational network

MicroRNA target sites were downloaded from MiRTarBase (Release 7.0, Chou *et al*., 2018) and converted into a compatible input format using mirTarbase_convert.py. Output files from microME.py were combined with mirTarbase_convert.py output files to produce an edge list of regulators to target interactions. Duplicate interactions were collapsed using collapse_maps.py and the number of interactions recorded as an edge weight.

### Mean filtering of TFBS to TSS interactions

For each dataset the number of binding sites for each TF was recorded at each TSS. Following the removal of outliers (> 2 SD of the mean), the mean number of binding sites for each TF is calculated. If the average number of binding sites at TSSs for a given TF is n, then any TSS where fewer than n sites are observed are removed from the network. Means for interactions are calculated after removing extreme outliers. TSS, where ≥n sites are found, would be maintained.

Filtering is performed by edge_weight_filter.py and run on the collapsed network generated in the previous step.

### Network numbers

The number and type of edges in the networks were counted using the count_edge_type.py script. This cycles over the network and quantifies the type of interactions seen in the network. e.g. protein_coding-microRNA, microRNA-protein_coding and protein_coding-protein_coding.

### Motif identification

Network motifs were identified by looking for patterns within the transcriptional-translational network edge list utilising the custom python script get_motifs_quicker.py. Autoregulatory loops are calculated first as other motifs are dependent on these loops. TFs in autoregulatory loops were then used to shortlist the search for further motifs. Motif discovery is based on boolean logic looking for distinct patterns on interaction. For example, for autoregulation instances were identified where gene-A targets gene-A. Motifs were identified by looking at all patterns of interaction required to generate the motif.

### Randomisation of Networks

Gene networks were randomised 1000 times utilising python script rewire.py by randomly swapping network edges. The swapping of edges was constrained such that microRNA to target interaction and TF to microRNA interactions were randomised separately. This prevents microRNAs being able to target the TSS of genes for example. Motifs were identified in each random network and compared to the real data. The number of edges exchanged was 1.1 x the number of edges per group (protein_coding, microRNA). Any edge swaps which resulted in the same edge or duplicated an edge already in the network were discarded.

Networks statistics were calculated as in Equation 1 and Equation 2 (North *et al*., 2003).

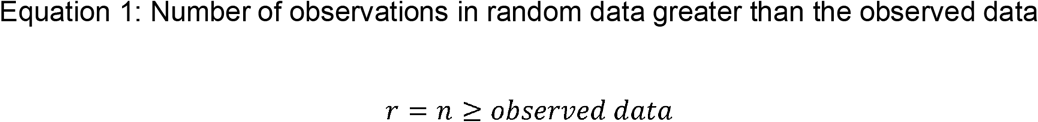

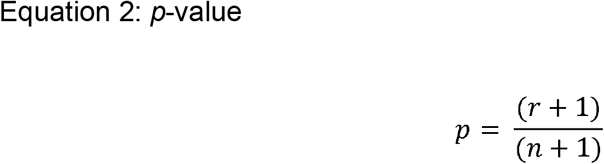

### Tissue networks

Tissue networks were generated by using the cell type and experiment data from the ReMap database and manually curating it into 28 tissue groups. Grouping was performed manually by identifying the tissue associated to each cell type. The network edges were then divided into network specific files based on the which tissue the data was grouped into.

Motifs were recalculated as above for each tissue network. The presence of each motif was then compared between each tissue network. The number of networks each TF was located in was calculated. If a TF was only found in one tissue network, then the data were discounted. For each motif a TF was located in was then compared between the tissue networks for which data for a given TF was found. Conservation was therefore the number of tissues where the motif was found as a percentage of the tissues where data for the TF exists.

## Supporting information

S1: Table of M1 motifs

S2: Table of M2 motifs

S3: Table of M3 motifs

S4: Table of M4 motifs

## Supplemental information

S1: Table of M1 motifs

S2: Table of M2 motifs

S3: Table of M3 motifs

S4: Table of M4 motifs

## Acknowledgements

The authors would like to acknowledge the assistance given by Research IT and the use of the Computational Shared Facility at The University of Manchester. The authors would like to thank Cerys Manning for proof-reading the manuscript.

## Third party data

TFBS and CRM data was obtained from ReMap2018 database (http://pedagogix-tagc.univ-mrs.fr/remap/). MicroRNA loci were obtained from miRbase (r21, http://www.mirbase.org/).

MicroRNA target data was obtained from miRTarBase (http://mirtarbase.mbc.nctu.edu.tw/php/index.php). Genome coordinates and TSS were downloaded from Ensembl (89) (http://www.ensembl.org/index.html).

## Author information

### Contributions

SGJ and NP supervised this study. TGM, SGJ and NP conceived and designed this study. All computational and bioinformatics experiments and subsequent analysis were conducted by TGM.

### Corresponding author

Nancy Papalopulu and Sam Griffiths-Jones

## Ethics declarations

### Ethics approval and consent to participate

Not applicable.

### Consent for publication

Not applicable.

